# The plasmid diversity of *Acinetobacter bereziniae* HPC229 provides clues on the ability of the species to thrive on both clinical and environmental habitats

**DOI:** 10.1101/710913

**Authors:** Marco Brovedan, Guillermo D. Repizo, Patricia Marchiaro, Alejandro M. Viale, Adriana Limansky

## Abstract

*Acinetobacter bereziniae* is an environmental microorganism with increasing clinical incidence, and may thus provide a model for a bacterial species bridging the gap between the environment and the clinical setting. *A. bereziniae* plasmids have been poorly studied, and their characterization could offer clues on the causes underlying the leap between these two radically different habitats. Here we characterized the whole plasmid content of *A. bereziniae* HPC229, a clinical strain previously reported to harbor a 44-kbp plasmid, pNDM229, conferring carbapenem and aminoglycoside resistance. We identified five extra plasmids in HPC229 ranging from 114 to 1.3 kbp, including pAbe229-114 (114 kbp) encoding a MOB_P111_ relaxase and carrying heavy metal resistance, a bacteriophage defense BREX system and four different toxin-antitoxin (TA) systems. Two other replicons, pAbe229-15 (15.4 kbp) and pAbe229-9 (9.1 kbp), both encoding MOB_Q1_ relaxases and also carrying TA systems, were found. The three latter plasmids contained *Acinetobacter* Rep_3 superfamily replication initiator protein genes. HPC229 also harbors two smaller plasmids, pAbe229-4 (4.4 kbp) and pAbe229-1 (1.3 kbp), the former bearing a ColE1-type replicon and a TA system, and the latter lacking known replication functions. Comparative sequence analyses against deposited *Acinetobacter* genomes indicated that the above five HPC229 plasmids were unique, although some regions were also present in other of these genomes. The transfer, replication, and adaptive modules in pAbe229-15, and the stability module in pAbe229-9, were bordered by sites potentially recognized by XerC/XerD site-specific tyrosine recombinases, thus suggesting a potential mechanism for their acquisition. The presence of Rep_3 and ColE1-based replication modules, different *mob* genes, distinct adaptive functions including resistance to heavy metal and other environmental stressors, as well as antimicrobial resistance genes, and a high content of XerC/XerD sites among HPC229 plasmids provide evidence of substantial links with bacterial species derived from both environmental and clinical habitats.

## Introduction

Plasmids are extra-chromosomal self-replicating DNA molecules that act as efficient vectors of horizontal gene transfer (HGT) across bacterial populations, thus facilitating adaptation of particular individuals and derived clonal lineages to novel and/or fluctuating environmental conditions [1,2]. Bacterial plasmids are assemblies of different modules encompassing replication, mobilization and stability functions in what is defined as the plasmid backbone [3]. This conserved structure is generally accompanied by accessory genes that provide adaptive functions, including pathogenicity/virulence determinants, antimicrobial resistance genes, and/or other traits depending on the selective context [1,3]. Plasmids are also carriers of mobile genetic elements such as insertion sequences (IS), transposons, and different genomic islands, thus becoming vessels for the transport of information from a communal gene pool [1]. They additionally provide scaffolds where different genetic rearrangements can occur via different homologous and non-homologous recombination processes, events that contribute not only to their own evolution but also the survival of their hosts under different evolutionary pressures [4].

The identification of plasmid types provides relevant information about their impact on the physiology of their hosts and their modes of transmission [3]. Current plasmid typing schemes exploit the more conserved backbone modules associated with replication (Rep typing) and/or mobility (MOB typing) [3,5]. Still, a caveat of these schemes is that plasmids lacking Rep or MOB genes escape classification schemes [6]. In this context, the rapid advance of whole genome sequence methodologies (WGS), by providing complete (or near-complete) plasmid sequences, has helped to overcome in part these limitations by providing valuable insights into the evolution of the accesory regions of the plasmids present in particular bacterial groups [7]. Thus, the characterization not only of the plasmid backbone but also of accesory genes including the complete repertoire of mobile genetic elements, and the identification of potential recombinatorial hot spots in the plasmid sequence, may provide many clues on the evolutionary processes that shaped their structures and drove their persistence in a given bacterial group, as well as their potentiality of dissemination to novel hosts [8-12].

We have recently demonstrated [13] that *A. baumannii* plasmids can exploit the XerC/XerD site-specific recombination system [14] to mediate fusions and resolutions of different replicons, some of them carrying *bla*_OXA-58_ and *aphA6* genes conferring carbapenem and aminoglycoside resistance, respectively. This may represent a powerful mechanism not only to drive plasmid structural rearrangements but also host range expansions [13]. Several authors have proposed in fact that the *Acinetobacter* XerC/XerD recombination system may mediate the mobilization of different DNA segments carrying adaptive or stability traits, a situation that includes antimicrobial resistance, heavy metal resistance, and toxin-antitoxin (TA) genes among others [6,13,15-18].

We have recently characterized by WGS a carbapenem-resistant *Acinetobacter bereziniae* strain isolated in Argentina designated HPC229 [19]. HPC229 carries a *bla*_NDM-1_-bearing carbapenem-resistance plasmid of 44 kbp designated pNDM229 [20], which displays a similar structure than plasmid pNDM-BJ01 carried by an *A. lwoffii* strain isolated in China [21]. This strongly suggested that pNDM229 was acquired by HPC229 as the result of HGT from other member of the genus, and selected due to the antimicrobial pressure common to the clinical setting [20]. As a species, *A. bereziniae* has a wide distribution and has been isolated from various sources including soils, vegetables, animals, and human-associated environments such as wastewaters [22-25]. More recently, *A. bereziniae* has been also increasingly associated to healthcare-associated infections in humans, and is currently considered an emerging *Acinetobacter* opportunistic pathogen [24,26]. In fact, although *A. bereziniae* isolates were in the past generally susceptible to most antimicrobials of clinical use [23], since 2010 a number of carbapenem-resistant clinical strains bearing IMP-, SIM-, VIM- or NDM-type metallo-β-lactamases (MβL) have been reported [20,24,27,28]. MβL genes are generally carried in these strains by plasmids, suggesting both the mechanism of their acquisition and potential spread to other Gram-negative species [20,24]. *A. bereziniae* might easily acquire plasmids, and has been proposed recently as a reservoir of clinically-relevant antimicrobial resistance genes [24]. Contrary to the acquired resistance plasmids mentioned above, the available information on the natural plasmids circulating among the population of this species and which may also contribute to the adaptation to different habitats is rather limited. This general lack of information on the *A. bereziniae* intrinsic plasmid repertoire reflects the case of most other species of the genus except for *A. baumannii* [29,30].

Here, we characterized in detail the whole plasmid content of the HPC229 strain, in an attempt to gain information on the intrinsic repertoire of *A. bereziniae* plasmids lacking antimicrobial resistance determinants and their putative role(s) in bridging the gap between environment and the clinical habitat. This included a comprehensive sequence analysis not only of the modules related to replication, stability, and mobilization, but also the identification of mobile elements, adaptive genes, and XerC/D recognition sites which may drive structural plasticity in their sequences and thus help in their persistence and the adaptation of their host(s) to different environments.

Parts of the results of this work were presented in the Symposium of Plasmid Biology 2018, Seattle, USA (Abstract 101).

## Materials and Methods

### HPC229 plasmids assembly and annotation

*Acinetobacter bereziniae* HPC229 is a carbapenem resistance clinical strain isolated from a blood sample of a 53-year-old female patient with leukemia [20]. The draft genome sequence of this strain was obtained using a 454 pyrosequencing platform (Roche Diagnostics) [19]. The revised assembled genome has been deposited at DDBJ/ENA/GenBank under the accession LKDJ02000000. Newly assembled genome sequences were annotated using the pipeline available at National Center for Biotechnology Information (NCBI).

Suspected plasmid sequences were analysed and assembled into putative replicons *in silico*, and the gaps inferred in the sequences were closed by PCR conducted on plasmid extracts with the aid of specifically designed primer pairs (S1 Table). The DNA sequences of all amplicons obtained in these assays were verified at the Sequencing Facility of Maine University. The overall analyses indicated the presence, besides of pNDM229, of five additional plasmids lacking antimicrobial resistance determinants (Table 1). The ORFs predicted by the NCBI pipeline in these plasmids were compared with the protein sequences deposited in the GenBank database using BlastP [31]. The CGview tool with default parameters (http://cgview.ca/) [32] was used for the visualization of GC content and GC skew.

**Table 1.**
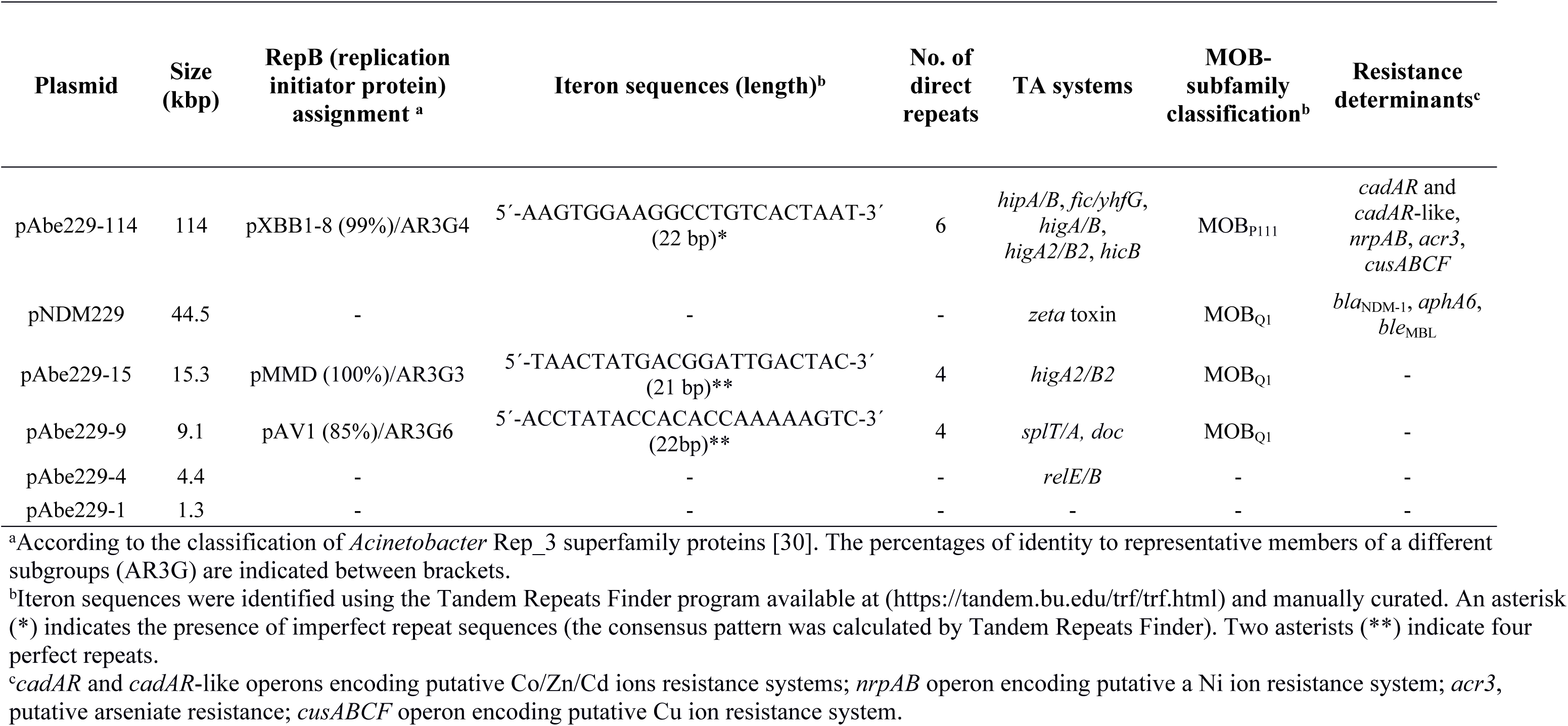
Characteristics of *A. bereziniae* HPC229 plasmids

### Plasmid isolation and S1 analysis

HPC229 plasmids were extracted using the Wizard DNA purification kit (Promega, Madison, WI, USA), and analyzed by 0.7% agarose gel electrophoresis and ethidium bromide staining following described procedures [13]. S1 nuclease treatment of plasmid extracts (S1 Fig) was conducted essentially as described previously [13].

### Comparative sequence analyses

The presence of mobile genetic elements among HPC229 plasmids was inspected using BlastN [31]. Only hits equal or higher than 70% nucleotide identity and minimum alignment lengths of 1,000 bp for pAbe229-114 or 300 bp for pAbe229-15, pAbe229-9 and pAbe229-4 were considered for further analysis. Graphical displays of nucleotide homologies between HPC229 plasmids with other *A. bereziniae* genomes were obtained using CGview (http://cgview.ca/) [32] employing a BlastN-homology search with an E-value cut-off of 1e-15.

### Bioinformatic identification of diverse genetic elements

Classification of membrane transport proteins was done using the Transporter Classification Database program (http://www.tcdb.org/) [33]. The screening for type II TA systems was conducted using TADB (http://bioinfo-mml.sjtu.edu.cn/TADB/) [34] and RASTA-Bacteria (http://genoweb1.irisa.fr/duals/RASTA-Bacteria/) [35] web-based search tools. Data retrieved from these databases were manually curated and complemented with BlastP-homology searches against the NCBI Protein database [31].

Insertion sequences (IS) were detected using IS Finder (https://www-is.biotoul.fr/) [36] and ISSaga [37]. The designation of novel IS elements were provided by the curators of the IS database [36], while transposon (Tn) designations were assigned by the Tn Number Registry [38]. The Tandem Repeats Finder program [39] was used to identify direct repeats by using default parameters. The GenSkew program (http://genskew.csb.univie.ac.at/) was used to predict global minima skew values generally associated to plasmid origins of replication [40].

### Strain assignation based on ANI calculations

Average nucleotide identity values (ANI) [41] were calculated and used for *Acinetobacter* species assignation. ANI were calculated from the published draft genome sequence data of the strains under study using orthoANI [42]. Briefly, each genome pair consisting in a query genome *versus* the reference strain CIP70.12 genome were first split into 1,020 bp fragments, which were used to run bidirectional BlastN searches (cut-off value higher than 70%). Only orthologous fragment pairs satisfying this requirement [42] were then taken into consideration for calculating ANI. An ANI cut-off value higher than 95% was adopted to define a given *Acinetobacter* species [43]. The accuracy of the species assignment by this procedure was evaluated by calculating the ANI corresponding to the non-*A. bereziniae* strains *A. guillouiae* CIP 63.46 and *A. gerneri* DSM 14967, which represent two *Acinetobacter* species showing the closest phylogenetic affiliation to *A. bereziniae* [22,25].

### ***A.*** bereziniae plasmid classification based on the comparison of Rep proteins

*A. bereziniae* plasmids bearing *rep* replication initiator protein (Rep) genes were classified following the scheme recently proposed by Salto et al. [30] for *Acinetobacter* plasmids. First, a search for known plasmid Rep domains as judged by the NCBI conserved domain database [44] was conducted among the *A. bereziniae* plasmid sequences reported in this work and those retrieved from *A. bereziniae* genomes available in the GenBank public database. Putative Rep candidates were then used as queries for a BlastP search against a local protein database that included all the Rep sequences reported by Salto et al. [30], and subsequently assigned to the corresponding *Acinetobacter* AR3G groups following the criterium described by these authors.

### Classification of A. bereziniae plasmids encoding MOB proteins

*A. bereziniae* plasmids bearing relaxase genes were classified following the scheme of MOB classification in families and subfamilies essentially following described procedures [30]. First, a local database that included all the MOB proteins used for the classification of *Acinetobacter* relaxases into MOB groups and subgroups was constructed [30]. Then, a search for proteins encoded in *A. bereziniae* genomic sequences available in the GenBank database with conjugation and/or mobilization functions was performed. This search retrieved 12 predicted protein sequences encoding for putative relaxases as judged by their homologies to described relaxase domains (NCBI conserved domain database) [44]. Among these 12 sequences, eleven showed the pfam03389 MobA/MobL domain and the other (encoded in pAbe229-114 and designated TraI*) showed the pfam03432 Relaxase/Mobilization nuclease domain. The eleven proteins bearing the pfam03389 domain were subjected to further phlylogenetic analysis. For this purpose, their N-terminal domains (first 300 amino acid residues) were first aligned with the MOB proteins of local database (see above) using MEGA6.06 [45] by employing ClustalW [46] with default parameters, and a Maximum-Likelihood (ML) phylogenetic tree was then generated using these alignments. To determine the best-fit protein substitution model, the tool included in MEGA6.06 was employed. This resulted in the use of the LG+G+I substitution model, taking into account the Akaike information criterion (AIC). Bootstrap values (100 replications) were also calculated using MEGA6.06 [45].

In the case of the TraI* relaxase encoded in pAbe229-114 (see above), the assignment was done using the best-hit match in a BlastP search against the relaxase local database.

### Search for XerC/D recombinases recognition sites in A. bereziniae plasmids

HPC229 plasmid sequences were first queried using Fuzznuc (http://www.bioinformatics.nl/cgi-bin/emboss/fuzznuc http://www.bioinformatics.nl/cgi-bin/emboss/fuzznuc) with default parameters with an ambiguous XerC/D nucleotide recognition sequence (NNTNYKYATAANNNNYWTTATSTKAWNN, whereY=C/T, K=G/T, W=A/T, S=G/C, N=A/T/C/G), inferred from the consensus 28-mer XerC/D recognition sequence determined for *A. baumannii* plasmids [13]. This search found 9 XerC/D recognition sites among these plasmids, which were complemented with 3 additional sites detected after a thoughtful visual inspection of plasmid sequences.

XerC/D recognition sites bearing a 6 nt-central region (cr6-XerC/D) detected in HPC229 plasmids were then used to infer a (degenerate) consensus XerC/D recognition sequence: (DHWYCKHATAANNNNNNTTATGTTAADT; where D=A/G/T, H=A/C/T), and used as query to identify equivalent sites in *A. bereziniae* sequences. A logo depicting the *A. bereziniae* XerC/D site frequency plot was then constructed using this information with the tool available at https://weblogo.berkeley.edu/logo.cgi.

### Ethics statement

This study involved sequence analysis of *Acinetobacter bereziniae* plasmids and did not implicate human specimens or participants.

## Results and Discussion

### Analysis of plasmids harbored by A. bereziniae HPC229

The different contigs obtained after HPC229 genomic pyrosequencing were extensively analyzed to search for plasmid sequences other than pNDM229 characterized in detail previously [20]. Among them we identified 5 putative plasmid sequences, which were thoughtfully assembled *in silico* and the resulting inferred final structures subsequently validated by PCR using specifically designed primer pairs (S1 Table, Figs 1, 2 and 4). Therefore HPC229 harbors, besides pNDM229, five other plasmids of approximate sizes of 114, 15.4, 9.1, 4.4, and 1.3 kbp which will be hereafter designated pAbe229-114, pAbe229-15, pAbe229-9, pAbe229-4, and pAbe229-1, respectively (Table 1; GenBank accession numbers are provided in S1 Table). Conventional plasmid extraction combined with S1 nuclease treatment and agarose gel electrophoresis analysis of the digested material (S1 Fig) indicated the presence of four DNA bands of approximate sizes of 15, 9, 4 and 1.3 kbp, additionally supporting the presence of pAbe229-15, pAbe229-9, pAbe229-4, and pAbe229-1 in these cells. The larger plasmids were not visualized by this procedure, a situation thay may have resulted from their much larger sizes and/or copy number.

### pAbe229-114 encodes five heavy metal resistance operons and a phage resistance system

The knowledge on *Acinetobacter* non-*baumannii* plasmids in general or *A. bereziniae* plasmids in particular, the functions they carry, and their transferability potential, is still scarce. This prompted us to conduce a detailed analysis of the backbone structure and accessory regions of the HPC229 plasmids detailed above.

pAbe229-114, the largest plasmid found in these cells, contains 114,007 bp, displays a GC content of 42.1% (Fig 1), and contains 111 ORFs from which 84 (i. e., 76%) encode proteins exhibiting known functions (S2 Table). It shows an *oriV-repB* replication region, with an encoded RepB replication initiator protein of the Rep_3 superfamily (pfam01051 conserved domain) displaying complete amino acid identity with their homologs encoded in *A. ursingii* UMB1319 and in *A. nosocomialis* 2010S01-197 (WP_043972782.1; S2 Table). pAbe229-114 was assigned to the *Acinetobacter* AR3G4 plasmid group, following the classification of *Acinetobacter* plasmids based on Rep_3 proteins recently proposed [30] (S3 Table). The *oriV* region, which is located 549 bp upstream of the *repB* start codon, contains six imperfect direct repeats of 22 bp each (Table 1, Fig 1) likely corresponding to iteron sequences playing roles during the initiation of replication and plasmid copy number control [47]. Comparative sequence analyses of the whole replication module including *repB* and the iterons indicated 99% nucleotide identity with a similar region located in plasmid pXBB1-8 of *A. johnsonii* XBB1 recovered from hospital sewage (Table 1) [48].

**Fig 1.**
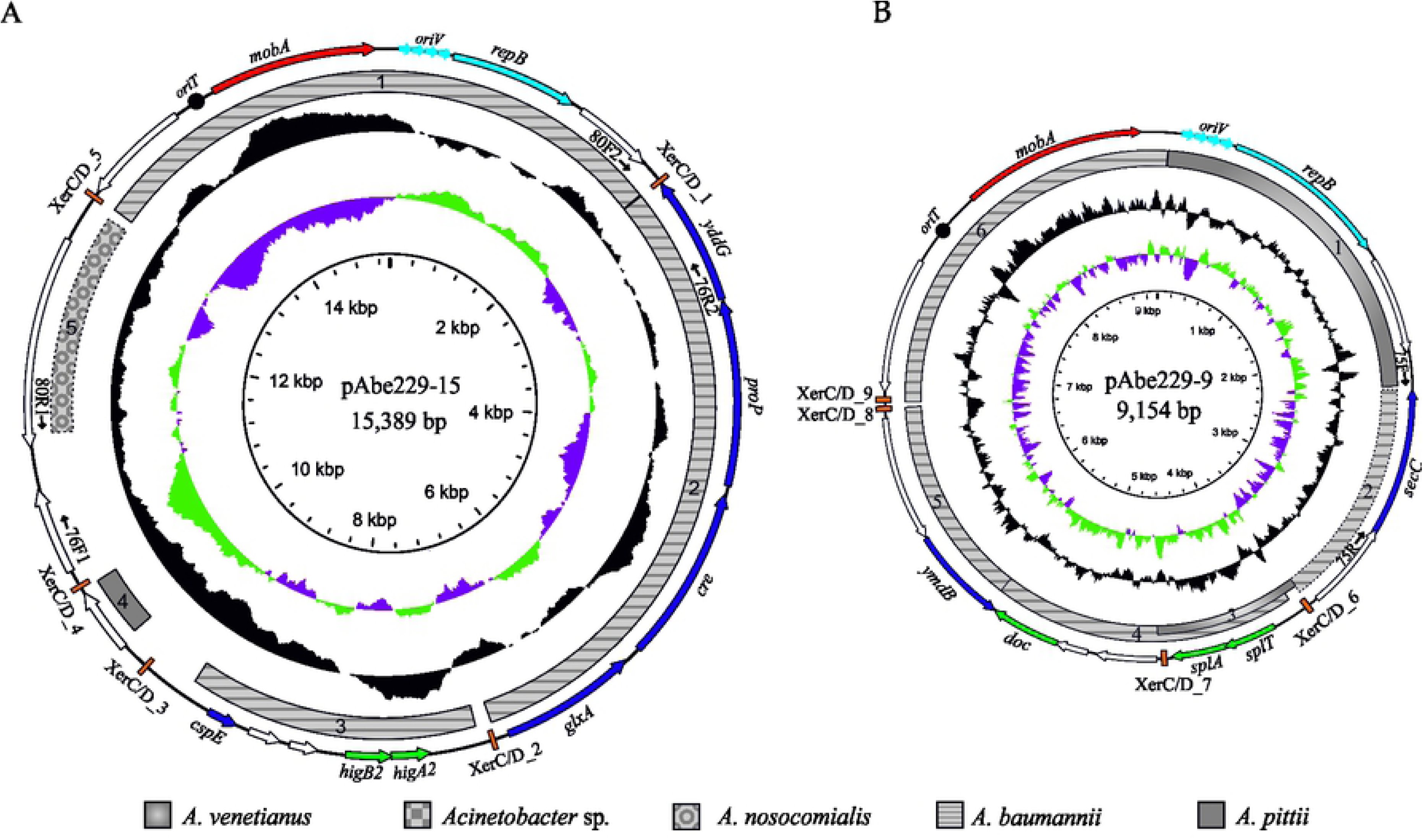
Schematic representation of pAbe229-114 plasmid. ORFs are shown as arrows indicating the direction of transcription. Disrupted/incomplete genes are indicated with a “Δ” symbol preceding the gene denomination. The six consecutive blue light arrows upstream of the *repB* gene denote the predicted iteron sequences linked to the *oriV* region (see Table 1 for details). The putative *oriT* located within the conjugal/transfer region is indicated with a closed circle. The Tn*6637* transposon bracketed by IS*Aba1* elements (one of them truncated, ΔIS*Aba1*) is highlighted in light yellow. The 9-bp (AATAAAGAT) direct repeats found at the insertion target site (DR) are indicated next to the external inverted repeats. From the outer circle inward, the circles display: i) the predicted ORFs. The colored arrows describe the location, identification, and direction of transcription of genes with described functions in databases. ORFs with undescribed functions are indicated by open arrows, ii) in grey with different designs, homologous regions described in plasmids/chromosomes of *Acinetobacter* non-*bereziniae* strains (see S4 Table for details), iii) GC content relative to the mean GC content of the plasmid, iv) GC skew, where green and purple represent positive and negative skew, respectively, v) scale in kbp. The hybridation sites of the PCR primer pairs used to verify the structure of pAbe229-114 (S1 Table) are also indicated.

In addition to the replicative module, pAbe229-114 encodes different stability functions, including ParA/ParB as well as HipA/HipB, Fic/YhfG, HigA2/HigB2, and HigA/HigB TA systems. For each of these TA systems, homologous were detected in other *Acinetobacter* species (Fig 1, S2 Table).

pAbe229-114 also contains a conjugative/transfer region (*traMLKJ*-Δ*traI*; S2 Table) in which the relaxase gene (pfam03432) is partially deleted at the 3’ region (hence the Δ*traI* designation). The translated protein (TraI*), however, still retains a complete relaxase domain. The conjugative/transfer region mentioned above showed 91% nucleotide identity with a homologous segment displaying a complete *traI* gene found in the conjugative plasmid pQKH54 (AM157767.1) from an uncharacterized bacterium [49]. A BlastP search using as query the N-terminal first 300 amino acids (relaxase domain) of TraI* showed 92% identity with a homologous relaxase corresponding to MOB_P111_ subfamily (Table 1) [30].

Comparative nucleotide sequence analysis of pAbe229-114 against *Acinetobacter* genomes deposited in databases allowed us to detect homology between several regions of this plasmid with other sequences present in other species of the genus (S4 Table; see the upper part of the Table for sequences present in *Acinetobacter* non*-bereziniae* genomes). For instance, the *parBA-repB* fragment encompassing both replication and stability functions (region 1, Fig 1, and S4 Table) showed more than 99% nucleotide identity with a homologous region carried by pAV3 from the environmental *A. venetianus* VE-C3 strain [50]. In addition, three of the pAbe229-114 TA systems, namely HipA/HipB, Fic/YhfG and HigA/HigB, were also identified in pAV3, with the corresponding DNA regions displaying sequence identities higher than 99% between them (regions 4 and 12, respectively, Fig 1, and S4 Table). Their presence likely fulfills an important role for plasmid stability in both replicons [51]. In turn, the HigA2/HigB2 TA system found in pAbe229-114 (region 9, Fig 1, and S4 Table) has probably been independently acquired from other sources, as inferred from the high percentage of identity between this region and a homologous sequence identified in *A. pittii*.

The region of pAbe229-114 comprising the *cadRA-like*-*feoAB* genes (cobalt-zinc-cadmium resistance genes, and ferrous iron transport operon, respectively) encoding heavy metal ions resistance functions shows 95% sequence identity with a homologous region located in plasmid pmZS of the *A. lwoffii* ZS207 environmental strain (region 2, Fig 1, and S4 Table). Moreover, the 6,091 bp-segment including the *cusFABC* genes encoding a RND efflux pump related to copper ion resistance showed 95% identity with a homologous region (region 3, Fig 1, and S4 Table) detected within the GI2-genomic island located on the chromosome of the *A. baumannii* LAC-4 clinical strain [52]. In turn, a 6,971 bp-DNA segment which includes the *cadAR* genes encoding cobalt-zinc-cadmium ions resistance contained in a truncated Tn*6018* element, and the *nrpAB* genes encoding nickel ion resistance [53,54],exhibited more than 98% nucleotide identity with pALWED3.1 of the *A. lwoffii* ED9-5a environmental strain (region 6, Fig 1, and S4 Table). Other adaptive regions probably involved in oxidative stress resistance encode functions responsible for the reduction of methionine sulfoxides in damaged proteins (regions 5 and 10, Fig 1, and S4 Table). These regions show homology to DNA sectors detected in *Acinetobacter* sp. WCHA45 and *A. johnsonii* plasmid pXBB1-9, respectively. The *fabAB* genes, which are located within region 5, have been related to intrinsic resistance to aminoglycosides [55,56]. Furthermore, a locus encoding a BREX system (S2 Table) reported to be involved in defense against bacteriophages [57], was detected in pAbe229-114 displaying higher than 92% identity with a homologous cluster present in plasmid pAV3 from *A. venetianus* VE-C3 (regions 7 and 8, Fig 1, and S4 Table). Remarkably, a region of this locus of around 1 kbp encompassing the central part of *pglX* (S4 Table) is not conserved between these plasmids. *pglX* encodes a protein containing an adenine-specific DNA methyltransferase motif, and variability in this gene has been documented and linked to a possible phase variation playing regulatory functions [57].

Of note, the overall observations above uncovered extensive regions of identity between the plasmid backbones of *A. bereziniae* pAbe229-114 and *A. venetianus* VE-C3 pAV3 (Fig 1, S4 Table). Comparative nucleotide sequence analyses between these two plasmids indicated in fact that they share around 50% equivalent sequences, including the replication and stability regions and part of the adaptive region. A further example is represented by region 4, in which these two plasmids share higher than 99% nucleotide identity (Fig 1, S4 Table). The additional presence of an integrase, two TA systems, as well as UmuC and UmuD error-prone DNA polymerase V subunit genes [58] in this region led us to conclude that this region constitutes a genomic island probably acquired by HGT. Besides the similarities between pAbe229-114 and other plasmids summarized above, unique regions were also identified in this plasmid including the conjugal transfer region as well as some segments involving genes contiguous to IS elements corresponding to the adaptive regions (Fig 1, S2 Table).

A search for mobile genetic elements in pAbe229-114 indicated the existence of 15 different IS (including complete IS and IS remnants) assigned to 7 different families (Table 2). The presence among them of IS*Aba12* and IS*Aba22* in multiple copies is consistent with previous observations indicating the ubiquitous distribution of these two IS among *Acinetobacter* genomes [59]. This suggested that pAbe229-114 has undergone several structural rearrangements, some resulting from IS insertions. Most of the IS shown in Table 2 have been reported previously in different members of the *Acinetobacter* genus [59]. However, we identified a novel IS element, which received the designation IS*Abe18* by the ISSaga database [36]. This IS*Abe18* is flanked by 9-bp direct repeats (Table 2), indicating a duplication at the insertion site. In turn, this not only suggest that it represents an active IS but also that it was recently acquired by the plasmid. It is worth noting that WGS sequence analysis indicated that all the above IS elements are missing in the *A. bereziniae* HPC229 chromosome (not shown), strongly suggesting that they were collected during transit of the plasmid through different bacterial hosts.

**Table 2.**
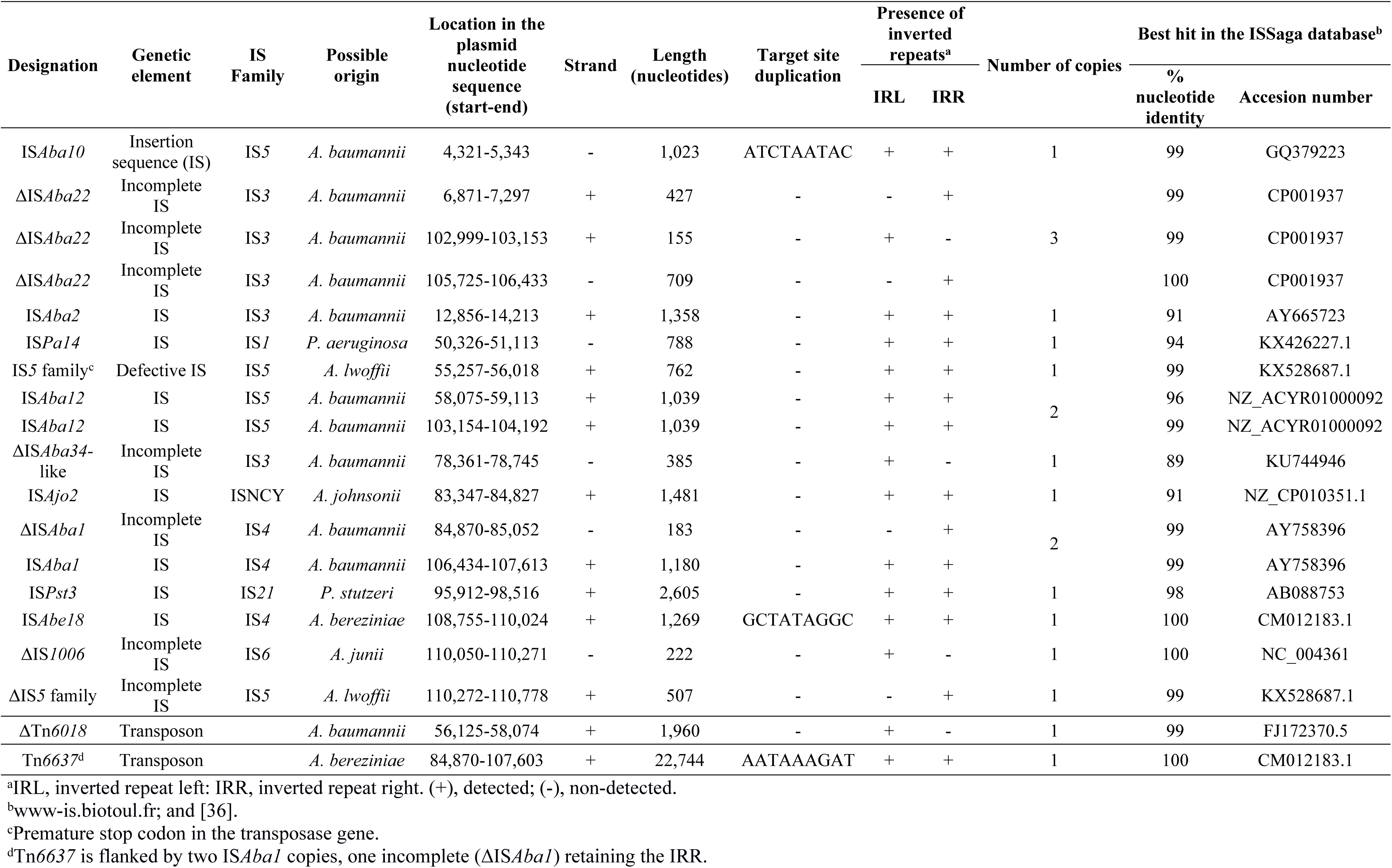
Mobile Genetic Elements detected in pAbe229-114

We also noted that pAbe229-114 bears a region of 18,283 bp (nucleotides 84,870-107,613 in Fig 1) bracketed by incomplete IS*Aba1* and IS*Aba22* elements, which exhibits a higher-than-average GC content (approximately 57%; see inner ring in Fig 1). When considering that the average GC content of *Acinetobacter* genomes is in average around 40% [60], it seems likely that this region was acquired by HGT from a donor from outside the *Acinetobacter* genus. In line with this assumption, a sequence exhibiting a similarly high GC content and displaying more than 85% nucleotide identity was identified in the draft genome sequences of *Pseudomonas stutzeri* B1 SMN1 (accession number AMVM00000000.1), an organism showing an analogous GC content [61]. Moreover, this region is contiguous in pAbe229-114 to a genetic element encompassing a *gshR* glutathione reductase gene (region 11, Fig 1, S4 Table) which is surrounded by an IS*Aba12* copy and a defective IS*Aba22* element that is immediately followed in turn by a complete IS*Aba1* copy (Fig 1, Table 2). IS*Aba1* exhibits a large impact on *A. baumannii* genomes [59], and therefore we hypothesize that the two IS*Aba1* elements bracketing the 22,744 bp region represent a composite transposon encompassing the oxidative stress resistance genes, the incomplete conjugal-transfer region, and the *gshR* gene. Supporting this notion, the two IS*Aba1* elements at the borders are limited by identical 9 bp-duplications immediately next to their external inverted-repeats, a hallmark of a recent IS*Aba1* transposition event [62]. This novel transposon was thus deposited at the Transposon Registry (https://www.lstmed.ac.uk/services/the-transposon-registry) under the designation Tn*6637* (Table 2). We also identified in pAbe229-114 an incomplete Tn*6018* transposon carrying cadmium and zinc resistance genes (Fig 1). Notably, the Tn*6018* element in pAbe229-114 lacks the *tnpA* transposase gene normally located downstream of the *lspA* gene [53], probably as the result of an IS*Aba12* insertion in this region. Downstream of the *cadR* gene, this defective transposon was additionally found to be interrumped by an IS*5* family element (Fig 1).

Overall the above analysis indicates a mosaic structure for pAbe229-114, most likely the outcome of the sequential insertion of different mobile elements accompanying the plasmid backbone followed by substantial rearrangements and deletions on the resulting structures. These events were most probably selected under varying external conditions during transit through different environmental and clinical bacterial hosts.

### The small HPC229 plasmids pAbe229-15 and pAbe229-9

*A. baumannii* clinical strains contain a varied repertoire of plasmids [2,3,6,13,17,29]. Many of them encode only a limited number of functions generally related to mobilization, a TA system, and a replicase of the Rep_3 superfamily, and are designated as “small” plasmids [6]. As detailed below, HPC229 plasmids pAbe229-15 and pAbe229-9 fulfill these traits, and could then be included in this category. pAbe229-15 (15,389 bp, 35% GC content; Fig 2A) contains 16 ORFs, 9 of them encoding proteins of known functions (S2 Table). Among them, replication (RepB), transfer (MobA), stability (a HigB2/HigA2 TA system), and adaptive (a YddG permease (DMT superfamily), a proline/glycine betaine ProP transporter, and a putative creatinase) functions were identified. In the replication module, the replication initiator protein RepB shows 100% amino acid identity with other Rep encoded in different *Acinetobacter* species (WP_012780181.1; S2 Table) including the Rep of pMMD carried by an *A. baumannii* clinical strain (YP_006961790.1; S3 Table) [16] and assigned to the AR3G3 group [30]. Four iterons of 21 bp were identified 44 bp upstream of the *repB* start codon (Table 1, Fig 2A), a region showing 100% nucleotide identity with a homologous region also located in pMMD. It is worth noting the presence in the adaptive region of pAbe229-15 of a gene encoding a protein showing both creatinase/prolidase (pfam01321) and creatine amidinohydrolase (pfam01321) domains. These two domains are commonly found in bacterial creatinases responsible for the hydrolysis of creatine to sarcosine and urea, thus providing carbon and nitrogen for growth [63]. The coexistence of genes encoding transporters and a transcriptional regulator belonging to the GlxA family in the same region (S2 Table) suggests that they shared a common metabolic pathway.

**Figure 2.**
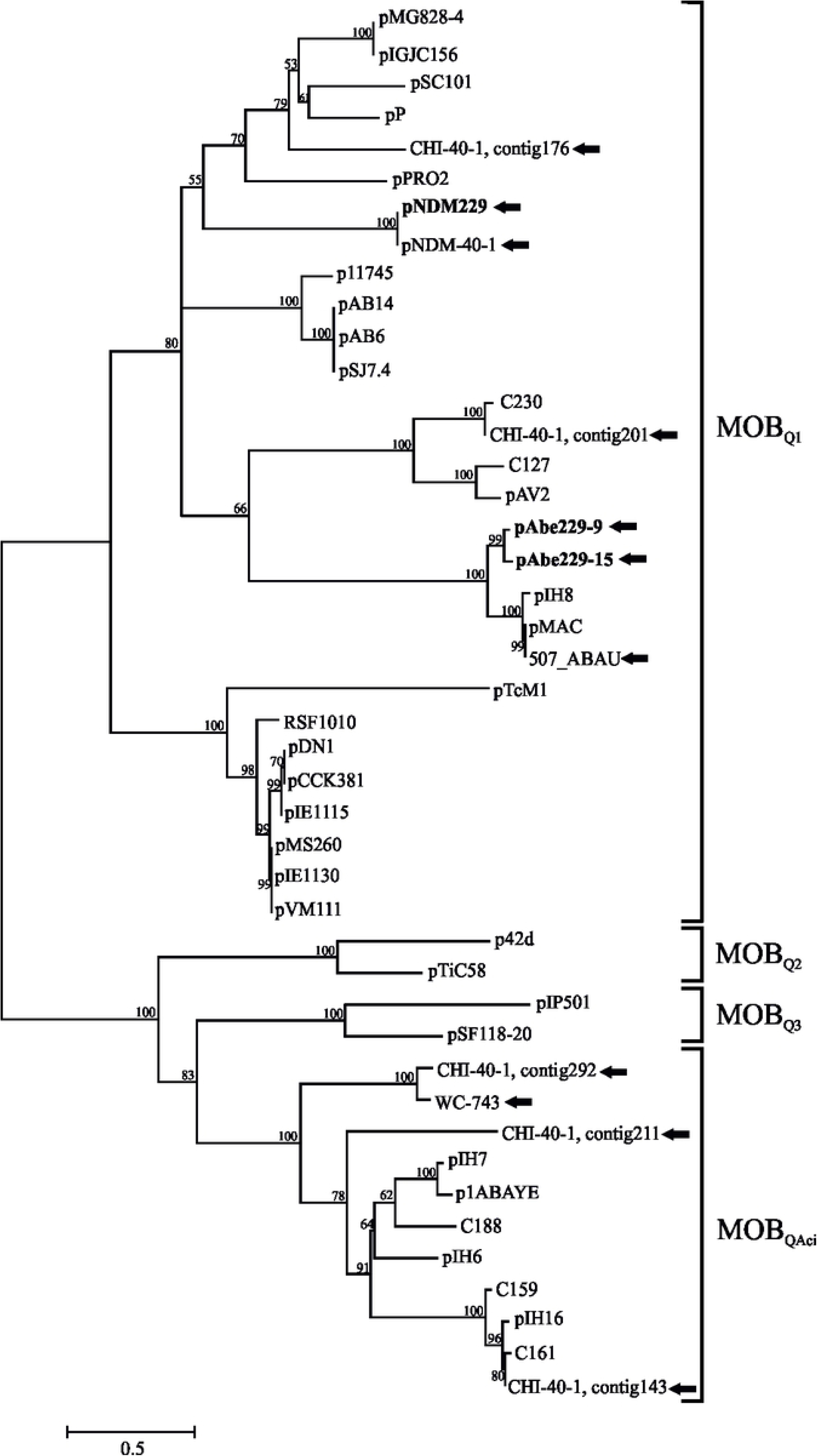
Schematic representation of plasmids pAbe229-15 (A) and pAbe229-9 (B). For details see the legend to Fig 1. Plasmids are not drawn to scale.

BlastN-searches uncovered significant sequence identities between different regions of pAbe229-15 with those of other *Acinetobacter* spp plasmids. For instance, the replication and transfer regions of pAbe229-15 showed 99% nucleotide identity with an equivalent region found in *A. baumannii* pMMD (region 1, Fig 2A, and S4 Table). In turn, the stability region including a TA system was homologous to an equivalent region from a plasmid found in *A. baumannii* AR_0052 (region 3, Fig 2A, and S4 Table). Homologous regions encoding similar adaptive functions were found between pAbe229-15 and a region of pABA-2f10 from *A. baumannii* ABNIH28 (region 2, Fig 2A, and S4 Table). Other segments of pAbe229-15 showed homology with regions of unknown functions found in *A. pittii* and *A. nosocomialis* (regions 4 and 5, respectively, Fig 2A). The overall analysis thus uncovered high levels of identity between different fragments of pAbe229-15 with equivalent regions found among members of the *A. calcoaceticus/A. baumanni* (ACB) complex.

pAbe229-9 (9,154 bp, 35.4% GC content; Fig 2B) contains 13 predicted ORFs, 7 of them coding for proteins of attributed functions (S2 Table). Its replication region contains a *repB* gene encoding a Rep_3 family protein with 100% amino acid identity with an homologous protein encoded in *A. pittii* PR331 (OTM21916.1) and 85% identity with an homologous protein encoded in pAV1 from the environmental strain *A. venetianus* VE-C3 (YP_001661463.1; S3 Table). Thus, similarly to the latter protein, pAbe229-9 RepB was assigned to the AR3G6 group following a recent classification [30]. The *oriV* region is composed by four perfect, directly oriented iterons located 56 bp upstream of *repB* (Table 1, Fig 2B). The whole replication region, including *repB* and the iterons, shows 79% nucleotide identity with a homologous segment of *A. venetianus* VE-C3 pAV1 (region 1, Fig 2B, and S4 Table). The stability region carries genes for a SplT/SplA TA system [64] and a toxin of the Doc-type. *splT*/*splA* homologous genes were identified also in pAV1 (region 3) and in pD36-4 of *A. baumannii* D36 (partial region 4, Fig 2B, and S4 Table). The latter plasmid also shares with pAbe229-9 a downstream segment (region 4, Fig 2B), which includes two ORFs of unknown function, and a *doc* gene encoding an orphan toxin of the Phd-Doc system (S2 Table). Concerning the transfer region, a 2,637 bp-homologous segment including a *mobA* gene was found in *A. baumannii* pMMD (region 6, Fig 2B, and S4 Table). Interestingly, the latter fragment also exhibits 93% nucleotide identity with a partial segment in region 1 of pAbe229-15 (Fig 2A) that spans from nucleotide positions 6,828 to 9,154 and continues from nucleotides 1 to 313 (Fig 2B). This strongly suggested that this region was exchanged in the past between these two HPC229 replicons.

### Phylogenetic analysis of the relaxases encoded by plasmids pAbe229-15 and pAbe229-9

The two *mobA* relaxases encoded in pAbe229-15 and pAbe229-9, respectively (S5 Table) were phylogenetically characterized following described procedures [30]. The corresponding N-terminal domain sequences were aligned with other 42 bacterial plasmid relaxases (S5 Table), and a ML phylogenetic tree was subsequently constructed (Fig 3). This analysis revealed that both relaxases clustered with members of the MOB_Q1_ subfamily, in particular within a subclade encompassing the relaxases from *A. baumannii* plasmids pMAC and pIH8. On the contrary, the relaxase encoded in pNDM229 [20] clustered within a different MOB_Q1_ subclade (Fig 3). It is worth remarking that in all HPC229 plasmids carrying *mobA* genes the genetic organization of the mobilization region is similar to that described for other *Acinetobacter* plasmids exhibiting a single *mob* gene [65].

**Figure 3.**
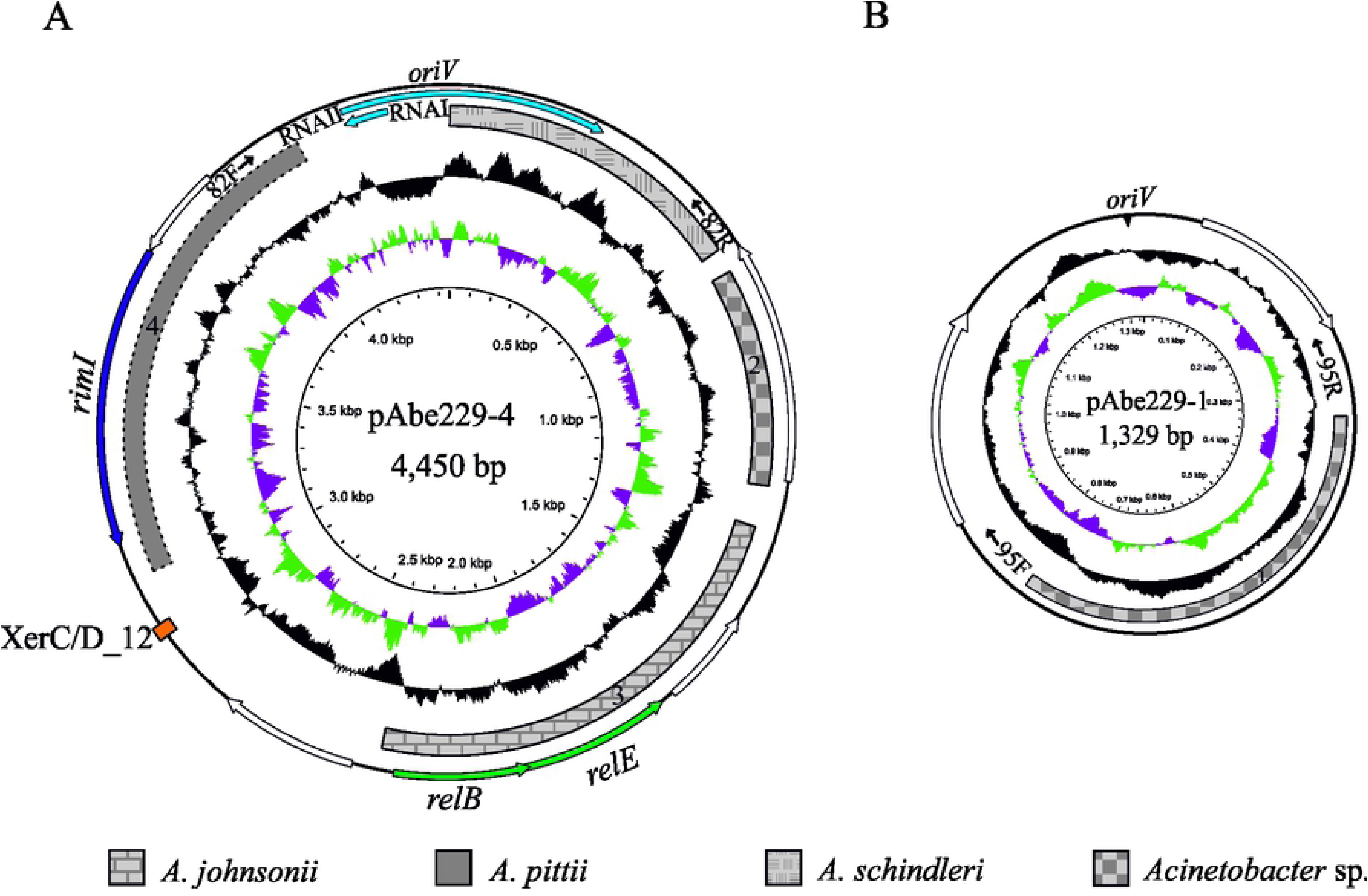
Phylogenetic analysis of HPC229 plasmid relaxases. A ML tree was inferred from the alignments of the first 300 amino acids of the N-terminal domains of MOB_Q_ relaxases from *A. bereziniae* and the *Acinetobacter* local database [3,30]. The MOB_Q1_, MOB_Q2_, MOB_Q3_ and MOB_QAci_ sub-families are indicated. Relaxases encoded in *A. bereziniae* genomes are specified with black arrows, and those corresponding to HPC229 plasmids are additionally highlighted in bold. In the case of the CHI-40-1 draft genome (GenBank accession number CDEL01000000.1), the contig number in which a given relaxase gene was identified is additionally indicated. The evolutionary scale (number of amino acid substitution per site) is indicated at the botton left. Bootstrap values higher than 50% (100 replications) are indicated at the branching nodes of the ML tree.

### The smallest HPC229 plasmids, pAbe229-4 and pAbe229-1, lack rep genes

The presence of small cryptic plasmids lacking known replication initiatior protein genes has been already described in *A. baumannii* [6]. Similarly, two of the plasmids found in *A. bereziniae* HPC229, pAbe229-4 (4,450 bp, 36.4% GC; Fig 4A) and pAbe229-1 (1,329 bp, 36.9% GC; Fig 4B) fall into this category (S2 Table). pAbe229-4 contains 7 ORFs, 3 of them with described functions including a RelE/RelBTA system and a *rimI* gene encoding a putative N-acetyltransferase. BlastN analysis disclosed significant regions of identity between pAbe229-4 with genomes of other *Acinetobacter* species, including the *relEB* locus which shows 87% nucleotide identity with a homologous region of plasmid pXBB1 of *A. johnsonii* XBB1 (region 3, Fig 4A, and S4 Table) and the *rimI* locus which shows 70% identity with a chromosomal region of *A. pittii* YMC2010/8/T346 (region 4, Fig 4A, and S4 Table).

**Figure 4.**
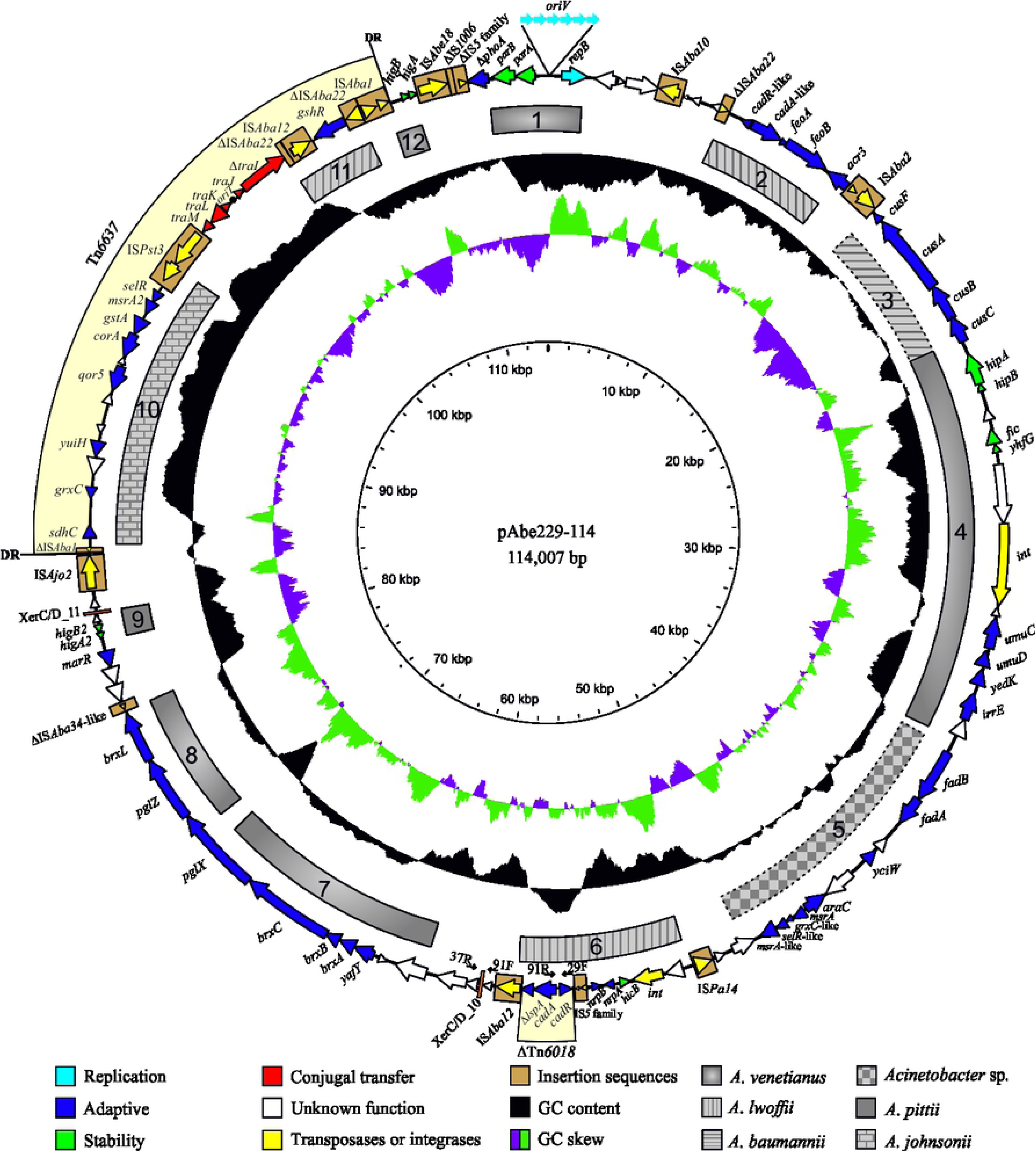
Schematic representation of plasmids pAbe229-4 (A) and pAbe229-1 (B). The regions encoding RNAI- and RNAII-homologous sequences in pAbe229-4 are highlighted in light blue, and the direction of transcription is also indicatedin each case. In pAbe229-1, the dark triangle denotes a high-AT region (putative *ori*V) predicted on the basis of the cumulative GCskew (see the text). Plasmids are not drawn to scale. For details, see the legend to Fig 1.

pAbe229-1, with only 1,329 bp (Fig 4B and S1 Fig), represents the smallest plasmid reported so far in an *Acinetobacter* genus member. It harbors 2 predicted ORFs with no homology in databases, but BlastN homology searches against the nucleotide GenBank database revealed significant homology to a 465 bp region of unknown function found in plasmid pM131-11 from an *Acinetobacter* sp. isolate (Fig 4B, S4 Table).

The absence of *rep* genes in pAbe229-4 and pAbe229-1 prompted us to search for alternate replication systems. In pAbe229-4 this analysis indicated the existence of sequences encoding homologous to RNAI (106 bp, between positions 4237 to 4342; Fig 4A) and RNAII (608 bp between position 4236 to 4450 and 1 to 393; Fig 4A), exhibiting 54% and 44% identity, respectively, to the equivalent regions described for ColE1 replicons [66,67]. pAbe229-4 would then represent, to our knowledge, the first example of a ColE1-type plasmid described in a species of the genus *Acinetobacter*. In the case of pAbe229-1, our analysis revealed only a high-AT region typical of a putative replication origin [68] (see potential *oriV* in the GC skew of Fig 4B), but its exact mechanism of replication is still unknown.

### Identification of XerC/D sites in HPC229 plasmids

Different authors [13,15-18] have noted the presence in different *Acinetobacter* plasmids of various modules, including antimicrobial resistance genes, heavy metal resistance genes, and TA systems among other genes, bordered by short DNA sequences recognized by XerC/XerD site-specific tyrosine recombinases (XerC/D sites). XerC/D sites are generally composed by two conserved motifs of 11 bp, separated by a less-conserved central region (cr) generally spanning 6 nt [69], although sites with cr regions of five, seven, and even more nucleotides have also been described [14,48,69,70]. The ubiquitous distribution of the XerC/XerD site-specif recombination system among bacteria has led to proposals that the XerC/D sites bordering these modules play roles in their mobilization and dissemination [13,15-18], although the exact mechanism is still obscure [17]. In this context, we have recently found [13] that at least some of the XerC/D sites located in a number of *A. baumannii* plasmids can in fact conform proficient pairs for site-specific recombination mediating the formation (and resolution) of plasmid co-integrates. We thus decided to investigate by bioinformatic procedures the abundance and location of XerC/D recognition sites in HPC229 plasmids (see Materials and Methods for details).

This approach allowed us to infer the existence of 12 putative XerC/D sites among 4 of the plasmids present in this strain: 5 in pAbe229-15; 4 in pAbe229-9; only 2 in pAbe229-114; and 1 in pAbe229-4 (Table 3). As previously observed for other bacterial XerC/D sites [13,16-18], the XerD recognition motif in HPC229 plasmids was more conserved than the XerC equivalent. In turn, the cr was the less conserved region both in sequence and in length (Table 3). Thus, while ten of the above 12 XerC/D sites show cr of an usual length of 6 nucleotides (cr6) displaying high sequence variability, one site (XerC/D_8, located in pAbe229-9) contained a cr of 5 nucleotides while other (XerC/D_5, located in pAbe229-15) a cr of 7 nucleotides in length (Table 3).

**Table 3.**
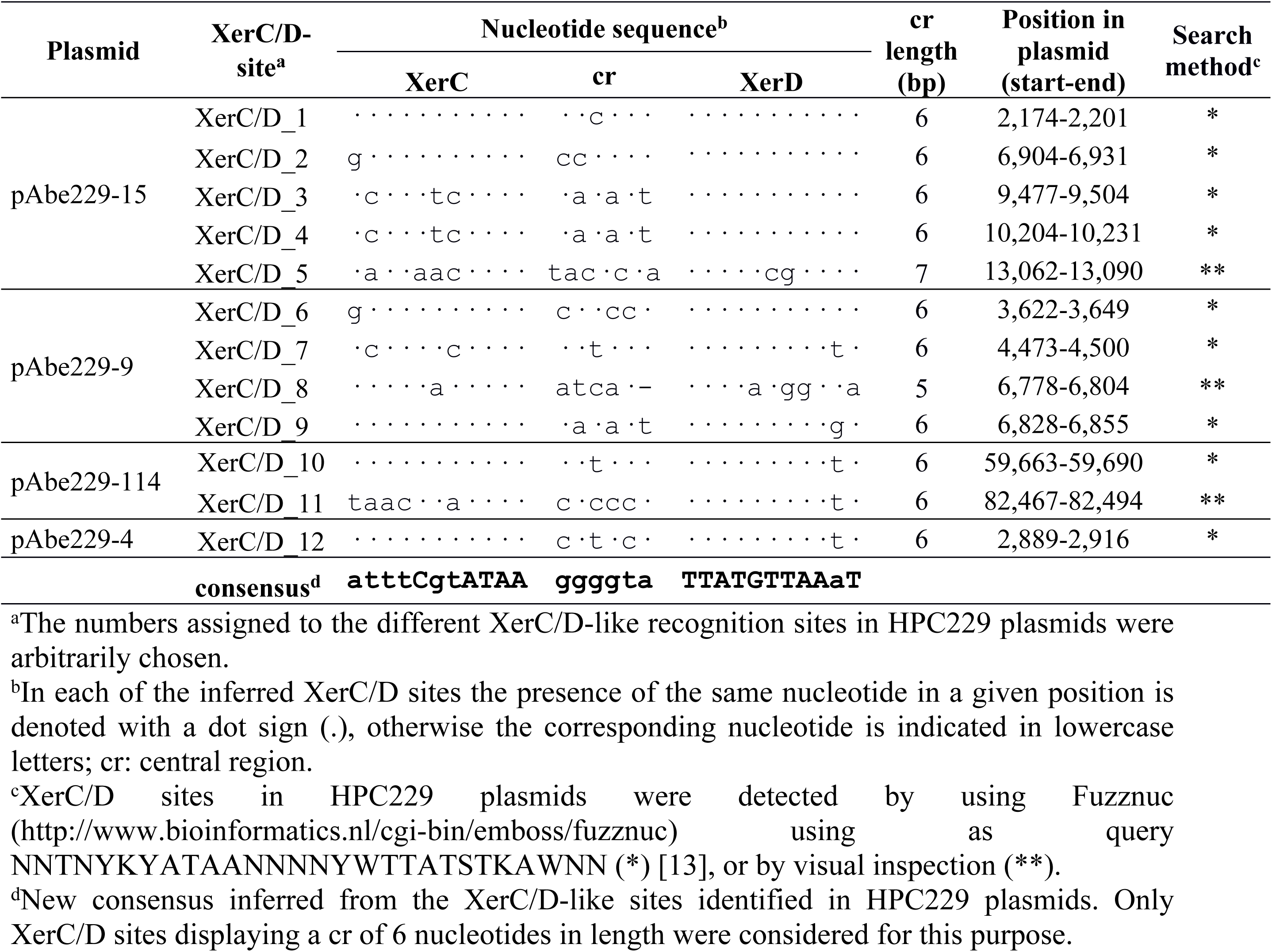
XerC/D sites in HPC229 plasmids

The above XerC/D sites were bracketing discrete regions in the corresponding HPC229 plasmid sequences (Figs 1 and 2). BlastN-homology searches using as query each of these regions indicated high levels of identity with similar regions carried by different *Acinetobacter* plasmids for some of them, including region 1 and 2 of pAbe229-15 (Fig 2A, S4 Table), and region 3 of pAbe229-9 (Fig 2B, S4 Table). It is worth noting also that a region of around 1 kbp in pAbe229-114, that includes the XerC/D_11 site and a downstream *higA2*/*higB2* TA system (region 9, Fig 1) showed high sequence identity with a homologous segment found in pOXA58-AP_882 of *A. pittii* AP_882 (S4 Table). This TA system is bracketed by a XerC/D pair in the latter plasmid (CP014479.1), thus suggesting that it may have been acquired by pAbe229-114 through XerC/D-mediated site-specific plasmid fusions followed by further plasmid rearrangements [13].

In summary, the presence of several XerC/D recognition sites in many of HPC229 plasmids, and the modular nature of the sequence regions that they border (see above) are consistent with the idea that XerC/D-mediated site-specific recombination events may have played relevant roles in the evolution of their structures [13].

### Analysis of plasmid idiosyncratic sequences (plasmid markers) in A. bereziniae genomes

To obtain further clues on the diversity of the plasmids housed by the *A. bereziniae* population, we conducted a comparative genomic analysis of genome sequences obtained from all strains assigned to this species available at GenBank (NCBI) database. For this purpose, we retrieved and analysed the available genomic sequence data corresponding to all seven strains classified as *A. bereziniae* in the database (S6 Table, top seven strains). We also conducted a BlastN search among the currently available *Acinetobacter* sp. genome sequences (GenBank-WGS database) using as query the *A. bereziniae* type strain CIP 70.12 *rpoB* gene sequence (GenBank accession number APQG01000052.1) to identify potential *A. berezeniae* strains not assigned to this species. Two strains showing *rpoB* sequences identities higher than 99% with CIP 70.12 *rpoB* emerged from this analysis, *Acinetobacter* sp. Ag2 and *Acinetobacter* sp. WC-743 (S6 Table), which prompted us to further delimitate their species assignations. We therefore calculated the percentage of average nucleotide identities (ANI) of these two strains when compared to the *A. bereziniae* type strain CIP 70.12 [22]. We also included in these comparisons other 6 strains previously assigned to *A. bereziniae* by other authors (S6 Table). As seen in this Table, ANI values higher than 98% were obtained for all of the 8 strains, largely exceeding in all cases the 95% cut-off value adopted for species assignation among the *Acinetobacter* genus [43]. In comparison, two *Acinetobacter* non-*bereziniae* strains that emerge closely-associated to *A. bereziniae* in phylogenetic trees such as *A. guillouiae* CIP 63.46 [22] and *A. gerneri* DSM 14967 [25] showed ANI values of 83 and 76%, respectively (S6 Table), thus validating the species assignations made above.

Plasmid contigs detected in the above *A. bereziniae* strains were first identified on the basis of the *rep* genes they encode. Hence, a list of Rep sequences was manually compiled based on the genomic annotation of all *A. bereziniae* strains under study (S6 Table), and in which proteins containing the chromosomal pfam00308 domain were excluded. Each of the identified Rep was used as query in a BlastP-homology search against a local protein database that included all the *Acinetobacter* replication initiator proteins of the Rep_3 superfamily (AR3) [30]. Sixteen out of the seventeen Rep proteins thus identified among *A. bereziniae* sequences fell into 11 of the 15 groups defined in this work [30], including AR3G1.1, AR3G1.2, AR3G1.4, AR3G2, AR3G3, AR3G5, AR3G8, AR3G9, AR3G11, AR3G12, AR3G13, AR3G14, and AR3G15 (S3 Table). A notable case was represented by *A. bereziniae* CHI-40-1, in which 7 Rep candidates were found distributed among 7 different AR3G groups (S3 Table). This supports the proposed existence of several different plasmids in this strain [28].

The remaining Rep candidate found in an *A. bereziniae* strain, KCTC 23199 (GenBank accession number WP_010591570.1; S3 Table), showed no significant identity to any of the above Rep_3 superfamily members. Moreover, it encompassed in the protein sequence an N-terminal replicase domain (pfam03090 family), a primase C-terminal 1 domain (pfam08708 family, PriCT_1) and a helix-turn-helix_28 domain (pfam13518 family). Further BlastP comparative searches against the GenBank protein database identified homologues also sharing these three domains not only among other *Acinetobacter* plasmids (S3 Table) but also enterobacterial plasmids including a *Klebsiella pneumoniae* plasmid (47% identity, 99% coverage, GenBank accession number SSI90041.1).

Two Rep proteins, encoded in *A. bereziniae* CIP 70.12 and KCTC 23199 sequences, were found to be identical to RepC (WP_000743064.1) codified in plasmid pAB3 of *A. baumannii* ATCC 17978 (S3 Table), which was assigned to AR3G14 [30]. However, these proteins contain a RepC (pfam06504) rather than a Rep_3 (pfam01051) superfamily domain, and were thus assigned to a novel group tentatively denominated ARCG1 (S3 Table). RepC superfamily members are found among IncQ-like group plasmids [71] where they function as iteron-binding *oriV* activators [72].

It follows that *A. bereziniae* is seemingly capable of hosting a wide variety of plasmids, the majority of them containing Rep_3 based replication modules. Still, the above observation strongly suggest that plasmids with other replication strategies are also replicative in this species.

We also analyzed the putative relaxases encoded in *A. bereziniae* sequences by undertaking the same phylogenetic approach used above for pAbe229-15 and pAbe229-9 (Fig 3, S5 Table). Eight proteins annotated as relaxases in *A. bereziniae* genomes were also included in this analysis with strain CHI-40-1 bearing 5 candidates, one of them in pNDM-40-1 (AHF22521.1; S5 Table) [28]. Our analysis (Fig 3) indicated that 4 out of these 8 proteins clustered within MOB_Q1_ together with the 3 relaxases detected among HPC229 plasmids, and the remnant 4 proteins clustered with members of the recently described MOB_QAci_ subfamily [30].

In summary, the wide repertoire of both replication and mobilization functions described above among *A. bereziniae* plasmids, and especially among strains HPC229 and CHI-40-1, reflect the plasmid diversity housed by this species.

### Comparative sequence analyses between HPC229 plasmids and other A. bereziniae genomes

We next analyzed whether the plasmids identified here in HPC229 share significant regions of identity with sequences of other *A. bereziniae* genomes (S6 Table, S1 Figure). We used for this purpose the draft genome sequences from 7 of the above indicated *A. bereziniae* strains, and excluded from the analysis strain XH901 since its reported genome contains only chromosomal sequences. This analysis indicated that the genomic island described in pAbe229-114 (region Abe4, S1A Fig) was also detected in the *A. bereziniae* strains NIPH 3, WC-743, CHI-40-1 and ABAU_507 (see also S4 Table; results obtained for sequences present in *A. bereziniae* genomes are shown in the lower part). The presence of this genomic island among these genomes suggests that it may have been acquired by HGT, although its presence in an *A. bereziniae* ancestor followed by differential losses in some lineages cannot be ruled out at this stage.

Other regions that deserve consideration in HPC229 plasmids are those encoding oxidative stress resistance mechanisms, including a region of Abe5 and the whole Abe10 region (S1A Fig). The corresponding genes were also identified in the CIP 70.12 and KCTC 23199 strains (S4 Table). Also, those regions encoding heavy metal (cadmium, cobalt, nickel and zinc) detoxification systems (region Abe6, S1A Fig) were also identified in strains NIPH 3, WC-743 and CHI-40-1 (S4 Table). It is worth noting that similar systems bearing strong adaptive significance have been described in plasmids from both pathogenic species of the genus such as *A. baumannii* [73,74] as well as from different environmental microbial populations [18,75]. Taken together, the above data reinforce the notion that these adaptive traits are being disseminated by plasmids among different bacterial populations.

Abe7 and Abe8 regions of pAbe229-114 (S1A Fig, S4 Table) encoding the type 1-BREX system (see above) were also found in sequences derived from the NIPH 3 and CHI-40-1 strains. In addition, a fragment including the *gshR* gene flanked by incomplete IS elements (Abe11, S1A Fig) was detected in CIP 70.12 and KCTC 23199 strains. Moreover, the Abe12 region bearing *higB/higA* genes is also present in CIP 70.12, KCTC 23199 and CHI-40-1 strains. Notably, the genes encoding all of these elements are contained in contig143 of strain CHI-40-1. This contig also harbors a *mobA* gene (Fig 3), thus suggesting a plasmid location of this TA system and supporting the above proposal that these systems are frequently mobilized by plasmids [76]. Altogether, the above comparisons indicate that *A. bereziniae* strains isolated in different geographical locations (S6 Table) share between them a significant number of DNA regions, most of them likely encoding adaptive functions. Still, pAbe229-114 carries a unique set of replication, stability and transfer genes as compared to other *A. bereziniae* strains (S4 Table). This suggests a high genetic plasticity for this species and the capability to mobilize DNA segments exhibiting adaptive functions among different *Acinetobacter* replicons.

Comparative sequence analyses done on pAbe229-15 showed homology to sequences present in other *A. bereziniae* genomes, with the exception of the region encompassing nucleotides 10,236-13,127 encoding two ORFs (orf13 and orf14) of unknown functions (S4 Table). This plasmid shares transfer and replication genes with 507_ABAU (region Abe1, S1B Fig), a creatine degradation pathway (region Abe2, S1B Fig) with WC-743, and the *higA2*/*higB2* genes with CHI-40-1 (part of region Abe3, S1 Fig). Moreover, the Abe3-4 and Abe4 regions flanked by XerC/D_3 and XerC/D_4 sites shows 98% sequence identity with an homologous region from *A. bereziniae* CHI-40-1 contig195 (S1B Fig, S4 Table). This segment in CHI-40-1 also encodes a Rep_3 member (CDEL01000195.1; S4 Table), thus suggesting that this contig corresponds to a plasmid. Thus, it is tempting to speculate that the Abe3-4 and Abe4 regions are shared by plasmids present in both CHI-40-1 and HPC229, and that are probably mobilized by recombination. Interestingly, the unique region of pAbe229-15 is flanked by sites XerC/D_4 and XerC/D_5 (S1B Fig), also suggesting an acquisition as the result of site-specific recombination.

In the case of pAbe229-9, a region displaying significant identity with sequences derived from strains CHI-40-1, WC-743, NIPH 3, and 507_ABAU (S1C Fig) carries the *splT/splA* genes (region Abe3; S4 Table). This region, also present in several *A. bereziniae* genomes, is bracketed by XerC/D_6 and XerC/D_7 sites. Moreover, the presence of a similar arrangement (including the bordering XerC/D sites) in pAV1 of *A. venetianus* VE-C3 also suggests a mechanism of acquisition of this TA encoding region mediated by site-specific recombination [13]. Abe6 homologous sequences probably encoding a transfer protein were also located in the 507_ABAU strain genome (S4 Table).

Regarding pAbe229-4, an homologous region of 2,352 bp including orf1, orf2, and the *relEB* TA system (Abe1, Abe1-2, Abe2, Abe2-3 and Abe-3, S1D Fig) was only detected in *A. bereziniae* Ag2 (S4 Table). Finally, pAbe229-1 homologous sequences could not be detected among *A. bereziniae* genomes other than HPC229.

Altogether, the above observations suggest that HPC229 plasmids are specific of *A. bereziniae* HPC229, since none of them showed significant coverage with sequences present in other *A. bereziniae* strains. Still, it is worth noting that 4 out of 5 HPC229 plasmids displayed homology with sequences present in strain CHI-40-1 including stretches of 42,386 bp (pAbe229-114), 2,571 bp (pAbe229-15), 1,422 bp (pAbe229-9), and 444 bp (pAbe229-4) (see total cover column, S4 Table). Furthermore, the modules conforming the HPC229 plasmid backbone are not widely represented in other members of this species, suggesting they were partially acquired from *A.* non-*bereziniae* species by HGT.

### XerC/D sites in A. bereziniae genomes

The presence of XerC/D recognition sites in *A. bereziniae* genomic sequences was also investigated. In principle we expected to detect the chromosomal cognate site (*dif*) involved in chromosome segregation [69], so we hypothesized that the additional XerC/D sites detected in a given genome should be carried by plasmids. To search for XerC/D sites among these sequences, a partially degenerate 28-nucleotides consensus query bearing a 6 nucleotides central region (cr), as inferred from HPC229 plasmid (Table 3), was used (see Material and Methods for details). Moreover, and given the lack of conservation in the cr on the consensus sequence mentioned above, we additionally modify the query by including a completely degenerate cr in this part of the sequence.

A total of 55 putative XerC/D sites were identified by this procedure in the eight *A. bereziniae* strains analyzed (S7 Table). Expectedly, only one XerC/D site was identified in strain XH901 lacking plasmid sequences (see above), probably reflecting the single chromosomal *dif* site. In most of the other strains the number of XerC/D sites varied between 2 and 7, with the exception of strains HPC229 (10 sites, Table 3) and CHI-40-1 (31 sites; S7 Table). This is in line with the above observations indicating that the latter two strains carry several plasmids, and suggests a correlation between strains adapted to the hospital environment and the presence of multiple XerC/D sites in the plasmids they carry [13].

With the information obtained above from all the 67 *A. bereziniae* XerC/D sites, we constructed a atttCgcATAAggggtaTTATGTTAAaT XerC/D consensus recognition site (S7 Table, see also Table 3). This consensus resulted very similar to that reported previously for the plasmids present in the *A. baumannii* Ab242 clinical strain (atTtcgtATAAggtgtaTTATgTtAaat) [13] with the exception of two mismatches, one at the XerC site and another at the cr. This opens the possibility of XerC/D-mediated site-specific recombinatorial events between plasmids of both species if they eventually come in contact within the same bacterial host.

## Conclusions

A detailed characterization of the *Acinetobacter* accessory genome can provide clues on the evolutionary dynamics of the members of this genus. The study of the “pan-plasmidome” in particular, including the whole set of plasmids harbored by pathogenic and environmental *Acinetobacter* strains, may help us to understand how these mobile elements actively mediate genetic exchanges between clinical and non-clinical habitats. In pursuit of this understanding, we analyzed here the plasmid content of *A. bereziniae* HPC229. This clinical strain represents, to our knowledge, the first *A. bereziniae* strain in which a complete and diverse pool of plasmids has been characterized. The ability of *A. bereziniae* HPC229 to harbor six plasmids, including pNDM229 carrying *bla*_NDM-1_ and *aphA6* resistance genes most likely acquired in the clinical setting [20] as well as the other plasmids characterized here and most probably derived from environmental sources, provides this strain with a great plasticity and the ability to thrive on different niches and under various selective pressures.

HPC229 plasmids pAbe229-114, pAbe229-15 and pAbe229-9 all encode replication initiator proteins of the Rep_3 superfamily similarly to the case of most other *Acinetobacter* plasmids [6,13,30,77]. This certainly opens the possibility for these three plasmids to replicate in other species of the *Acinetobacter* genus. On the contrary, pAbe229-4 and pAbe229-1 plasmids lack replication initiator protein genes. However, while the pAbe229-4 plasmid may use a ColE1-type replicon as suggested by sequence analysis, the replication mechanism of pAbe229-1 remains obscure.

The overall analysis suggest that *A. bereziniae* HPC229 plasmids could be regarded as chimeras of diverse origins. An outstanding example is pAbe229-114, which shares extensive backbone sequence similarity to pAV3 from *A. venetianus* VE-C3 isolated from polluted waters [78]. This correlation was not totally unexpected, providing that *A. bereziniae* is also isolated from wastewater products of human activities [25]. Moreover, similarity to pAV3, pAbe229-114 also contains various systems involved in the detoxification of heavy metal ions, probably reflecting a common origin in bacteria thriving in polluted environments. However, the presence of a relative high number of IS and IS remnants, a novel composite transposon, a genomic island, and many other adaptive functions also suggest that pAbe229-114 has been modeled by different events that included the capture of mobile elements and the incorporation and loss of different modules by site-specific recombination, while transiting trought different hosts under diverse environmental and selective pressures.

The small HPC229 plasmids pAbe229-15 or pAbe229-9 show features which may contribute to their dissemination by HGT. These include MOB_Q1_ relaxases, which could be assisted by helper plasmids from different incompatibility groups and/or chromosomally-located *tra* functions [65]. Interestingly, these small plasmids do not carry mobile elements but are enriched in XerC/D-recognition sites bracketing specific regions, which could mediate the formation of cointegrates with other XerC/D-containing plasmids by site-specific recombination [13]. In this context, our analysis of *A. bereziniae* genome sequences suggests a correlation between strains adapted to the hospital environment such as HPC229 and CHI-40-1 [20,28], and the presence of multiple XerC/D sites in many of the plasmids they carry.

In summary, our studies highlight the plasmid diversity carried by *A. bereziniae*, an organism that, by acting as a bridge among the nonclinical and clinical habitats, may contribute to the evolution and expansion of a plasmidome that facilitates the adaptation of other *Acinetobacter* species to these radically distinct environments.

## Author Contributions

ASL, PM, MB, AMV, and GR conceived and designed the work. MB performed the experimental work. MB and GR conducted the bioinformatic analysis. MB, ASL, PM, GR, and AMV analyzed the data. ASL, AMV, GR, and MB wrote the manuscript. All authors read and approved the final manuscript.

## Acknowledgments

We are indebted to Dr. M. Espariz for his generous help in the calculation of ANI values of the *A. bereziniae* genomes, and Dr. M. Pistorio for providing us the sequences of *Acinetobacter* spp. MOB_Q_ relaxases. We also thank P. Stothard for his generous help with the CGView software and the curators of ISSaga team for curating the data and making them publicly available.

## Supporting information

**S1 Fig. S1 nuclease analysis of HPC229 plasmids.** Plasmids extracted from *A. bereziniae* HPC229 without (lane 1) or with S1 nuclease treatment (lane 2) were resolved by agarose gel electrophoresis. The linearized forms of pAbe229-15, pAbe229-9, pAbe229-4 and pAbe229-1 are highlighted by black triangles. The final positions of the size markers (EcoRI/HindIII-digested Lambda DNA) are shown at the left margin.

**S2 Fig. Identification of homologous sequences to HPC229 plasmids in other *Acinetobacter* genomes.** The colored arrows in the outer ring describe the location, identification, and direction of transcription of genes in the corresponding plasmids with described functions in databases. ORFs encoding for unknown functions are indicated by open arrows. In the subsequent rings, the regions of homology between the corresponding plasmid sequences and genome sequences of other *A. bereziniae* strains (indicated by arcs of different colors, see key near the upper right margin) as detected by BlastN searches are shown. The strains have been located from the highest (outer rings) to the lowest (inner rings) sequence coverage. The height of the colored arcs in each case is proportional to the percentage of nucleotide identity obtained in the BlastN search. BlastN hits with ≥70% nucleotide identity and a minimum alignment length of 1,000 bp (pAbe229-114) or 300 bp (pAbe229-15, pAbe229-9 and pAbe229-4) are detailed in S4 Table. The darker lines in the arcs mark overlapping hits. The external Abe1-Abe12 grey regions indicate homologous sequences found in *Acinetobacter* non-*bereziniae* genomes. (For details see Figs 1, 2, and 4; and also S4 Table).

S1 Table. Oligonucleotide primers used for PCR analysis.pdf

S2 Table. Genes identified in HPC229 plasmids.xlsx

S3 Table. *A. bereziniae* plasmid replicases classification.xlsx

S4 Table. Homologous regions in other *Acinetobacter* spp.xlsx

S5 Table. Plasmid relaxases used in this work.xlsx

S6 Table. Characteristics of *A. bereziniae*, *A. guillouiae* and *A. gerneri* reference strains.xlsx

S7 Table. XerC/XerD sites in *A. bereziniae* genomes.xlsx

